# A thermostable, closed, SARS-CoV-2 spike protein trimer

**DOI:** 10.1101/2020.06.15.152835

**Authors:** Xiaoli Xiong, Kun Qu, Katarzyna A. Ciazynska, Myra Hosmillo, Andrew P. Carter, Soraya Ebrahimi, Zunlong Ke, Sjors H.W. Scheres, Laura Bergamaschi, Guinevere L. Grice, Ying Zhang, The CITIID-NIHR COVID-19 BioResource Collaboration, James A. Nathan, Stephen Baker, Leo C. James, Helen E. Baxendale, Ian Goodfellow, Rainer Doffinger, John A.G. Briggs

## Abstract

The spike (S) protein of SARS-CoV-2 mediates receptor binding and cell entry and is the dominant target of the immune system. S exhibits substantial conformational flexibility. It transitions from closed to open conformations to expose its receptor binding site, and subsequently from prefusion to postfusion conformations to mediate fusion of viral and cellular membranes. S protein derivatives are components of vaccine candidates and diagnostic assays, as well as tools for research into the biology and immunology of SARS-CoV-2. Here we have designed mutations in S which allow production of thermostable, crosslinked, S protein trimers that are trapped in the closed, pre-fusion, state. We have determined the structures of crosslinked and non-crosslinked proteins, identifying two distinct closed conformations of the S trimer. We demonstrate that the designed, thermostable, closed S trimer can be used in serological assays. This protein has potential applications as a reagent for serology, virology and as an immunogen.

## Introduction

Severe acute respiratory syndrome coronavirus 2 (SARS-CoV-2) is the virus responsible for the coronavirus disease of 2019 (COVID-19) pandemic ^1,2^. SARS-CoV-2 belongs to the beta-coronavirus family and is closely related to SARS-CoV-1 which was responsible for the 2003 SARS epidemic ^3^. Similar to other coronaviruses, Spike (S) protein is the major glycoprotein on the SARS-CoV-2 virus surface. S protein is responsible for receptor binding and subsequent virus uptake and fusion with the target cell, and is a dominant target of the immune system ^4^. S protein is trimeric and has two distinct structural states – prefusion and postfusion ^4^. Recognition by the immune system of the prefusion state displayed on the virus surface is crucial to mount an effective immune response ^5^.

S protein trimers of SARS-CoV-1 and middle east respiratory syndrome (MERS)-CoV are unstable in their metastable prefusion state ^6,7^. Stimuli including protease digestion ^8,9^, receptor binding, and antibody binding ^8^, can promote structural transition of S into the postfusion state. SARS-CoV-2 S protein exhibits similar instability and S trimers on the surface of SARS-CoV-2 virion or solubilised by detergent undergo structural transition into the postfusion state during purification ^10,11^. Successful immunisation strategies require stable antigens, and instability of the prefusion state of enveloped virus fusion proteins presents a challenge for vaccine development. For HIV-1, introduction of the SOSIP mutations (a disulphide bond and a stabilizing proline) into HIV-1 GP160 ^12^, and for paramyxoviruses, introduction of disulphide bonds and cavity-filling mutations into the fusion protein F, were required to stabilise these proteins in their prefusion state for structural or vaccinology studies ^5,13^. Attempts have been made to stabilise SARS-CoV-1 and MERS-CoV S proteins in their prefusion state by introduction of 2 prolines within S2 to prevent central *α*-helix extension, a structural change needed for transition into the postfusion state ^6,7^. Here, we found that such stabilisation does not maintain coronavirus S proteins in the prefusion form for a prolonged period of time.

Cryo-electron microscopy (cryo-EM) studies have shown that the fusion core of the S proteins of SARS-CoV-1 and MERS-CoV is crowned by the receptor binding domain (RBD), surrounded by the N-terminal domains (NTDs) to suppress fusion ^5^. Both NTD and RBD exhibit some flexibility, but the major conformational dynamics are between a closed and an open conformation. In the closed conformation the RBD is bound in-trans into a pocket formed by NTD and RBD of the neighbouring monomer, and the receptor binding site is largely occluded. In the open conformation the RBD points upwards and the receptor binding site is exposed ^14,15^. Similar RBD conformational dynamics has been observed for SARS-CoV-1 and SARS-CoV-2 S protein ^9,16^. It has been suggested that these conformational dynamics are related to both receptor binding and subsequent fusogenic activity ^14,15,17^. Binding of receptor or antibody to the open conformation appears to stimulate transition of S into the postfusion state ^8,18^. In addition, RBD dynamics is modulated by protease digestion where S proteins are cleaved into S1 and S2 subunits, which also appears to stimulate transition to the postfusion state ^8,9^.

Based on this understanding of S protein fusion mechanism, we sought to stabilise the S protein in the prefusion state by trapping it in the closed form using engineered disulphide crosslinks. Our motivation for doing so was four-fold. First, for use in vaccinology – a stabilized trimer may elicit an improved immune response as part of a vaccine, and stabilized vaccines may be more easily distributed. Second, to induce different immune responses – a closed trimer can be expected to lead to production of a different antibody repertoire to that raised against open S proteins. A successful SARS-CoV-2 vaccine will likely need to induce a breadth of immune response, and a successful immunotherapy may also require a combination of antibodies. Third, for diagnostics – a more thermostable trimer could enable better diagnostics in regions with poor infrastructure, and closed trimer can be used to distinguish different immune responses in diagnostic assays. Understanding the functional relevance of different antibody responses is critical to establish meaningful assays for protective immunity. Fourth, for basic research including structural biology. Here we have designed an exclusively closed trimeric S protein which is stable to incubation at 60°C and to prolonged storage.

## Results and discussion

### Construct design

Based on published structures of trimeric S protein ^19^, we identified two amino acid pairs at an appropriate distance for disulphide bond formation, and in positions where we expected covalent crosslinking to stabilize the closed form of the trimer. These pairs, residues 383 and 985 (crosslink 1, x1) and residues 413 and 987 (crosslink 2, x2), are illustrated in **Figure 1a**. Starting with the previously published stabilized trimeric ectodomain constructs in which the S1/S2 cleavage site (residues 682-685) was replaced by GSAS and two stabilizing prolines were inserted at positions 986 and 987 (S-GSAS/PP) ^16^, we inserted cysteines at the identified positions to generate the constructs S-GSAS/PP/x1 and S-GSAS/PP/x2 (**Figure S1**). During preparation of this manuscript, two preprints have reported constructs similar or equivalent to S-GSAS/PP/x1 ^20,21^. In addition, we inserted cysteines at the same positions in a related construct in which the residues PRRA have been deleted at the S1/S2 cleavage site, to leave a single arginine resembling the cleavage site in SARS-CoV-1 and SARS-CoV-2 potential progenitor RaTG13 ^1^ spikes (R), generating S-R/PP/x1 and S-R/PP/x2 (**Figure S1**). Finally, anticipating that the proximity of the stabilizing prolines at positions 986 and 987 to the closed RBD might affect the RBD conformation, we additionally engineered x1 and x2 into the GSAS and R constructs without the stabilizing proline mutations, generating S-GSAS/x1, S-GSAS/x2, S-R/x1 and S-R/x2 (**Figure S1**).

**Figure 1:**
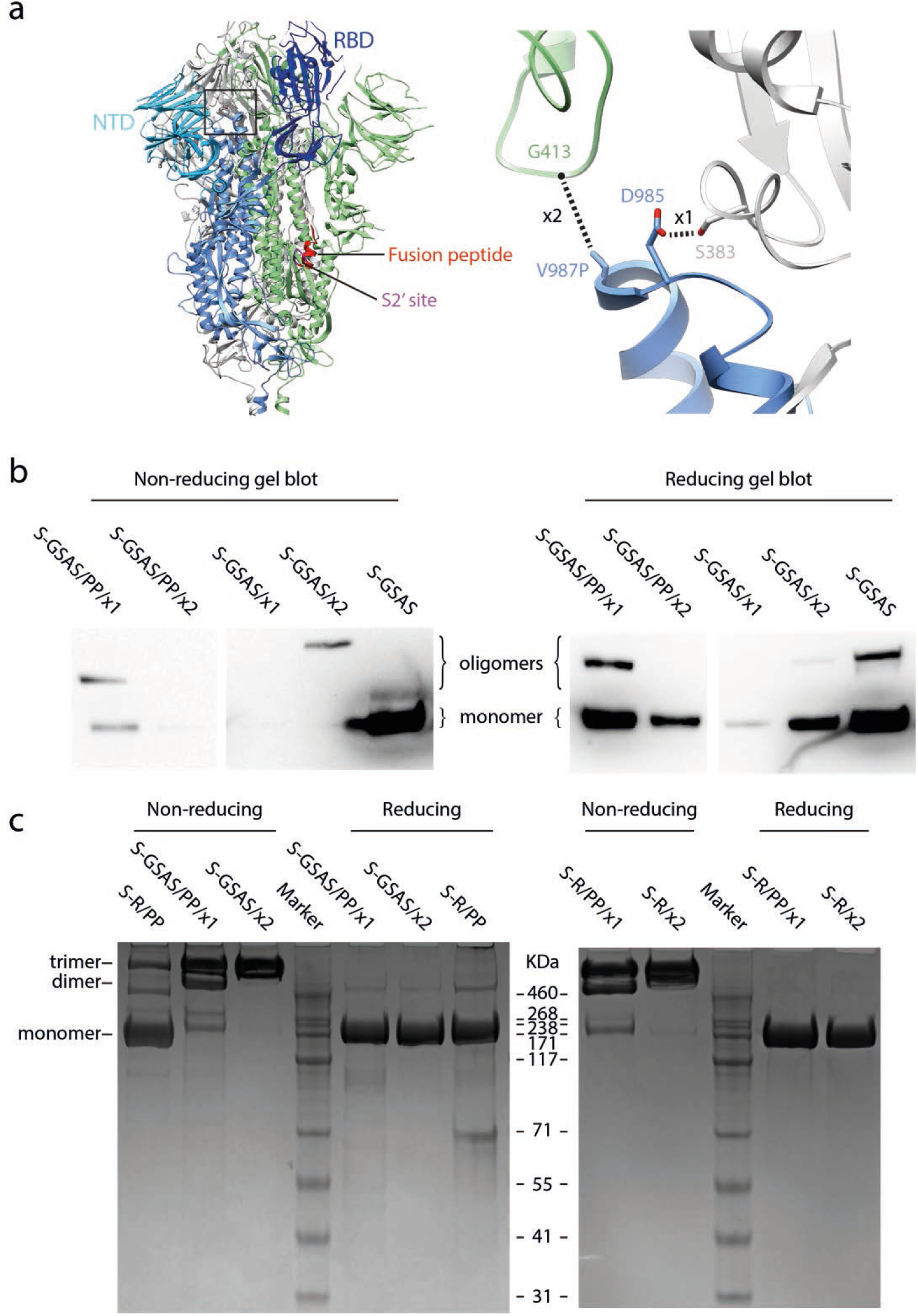
Design and expression of S protein constructs. **a)** A schematic of the structure of trimeric S in its closed, prefusion form (PDBID: 6VXX) illustrating the location of the RBD and the NTD, The right hand panel shows the positions of crosslinks x1 and x2. Note that in 6VXX and other published SARS-CoV-2 S protein structures, residues K986 and V987 have been mutated to proline. **b)** Western blots using antibodies against the C-terminal 6xHis tagged S-protein constructs in small-scale expression supernatants under non-reducing and reducing conditions. S-GSAS/PP/x2 and S-GSAS/x1 showed very low expression levels and were not considered further. **c)** Comparison of nonreducing and reducing Coomassie stained gels from medium-scale purifications of S-protein constructs. S-GSAS/PP/xl and S-R/PP/x1 show a mixture of trimers and lower order oligomers under non-reducing conditions, whereas S-GSAS/x2 and S-R/x2 show trimers under these conditions.

### Protein expression and crosslinking

Expression of all GSAS constructs in mammalian Expi293 cells was assessed by western blotting (**Figure 1b**). We were able to express S-GSAS/PP/x1 and S-GSAS/x2, but not S-GSAS/PP/x2 or S-GSAS/x1. Expression levels of the cysteine mutants are somewhat lower than those of the initial construct, possibly due to the effect of engineered cysteines on protein folding.

Based on the western blot results, we performed scaled-up expression and purification of S-GSAS/PP/x1 and S-GSAS/x2, as well as S-R/PP/x1 and S-R/x2, Purified proteins were assessed for crosslinking on non-reducing gels. S-GSAS/PP/x1 and S-R/PP/x1 have bands corresponding to trimers, dimers and monomers, while S-GSAS/x2 and S-R/x2 run exclusively as trimers (**Figure 1c**). Under reducing conditions only monomers were observed, confirming that the oligomeric states present in the non-reducing gel are due to disulphide cross-links.

### Stability of crosslinked mutants

We imaged the proteins by negative stain electron microscopy (EM) to assess their oligomeric states (**Figure 2, left column**). Freshly purified non-crosslinked variants, as well as the x1 crosslinking mutants, show a majority of trimers with some disintegrated oligomers. The x2 crosslinked mutants, which migrate exclusively as trimers in gels, show almost exclusively trimers by negative stain EM.

**Figure 2.**
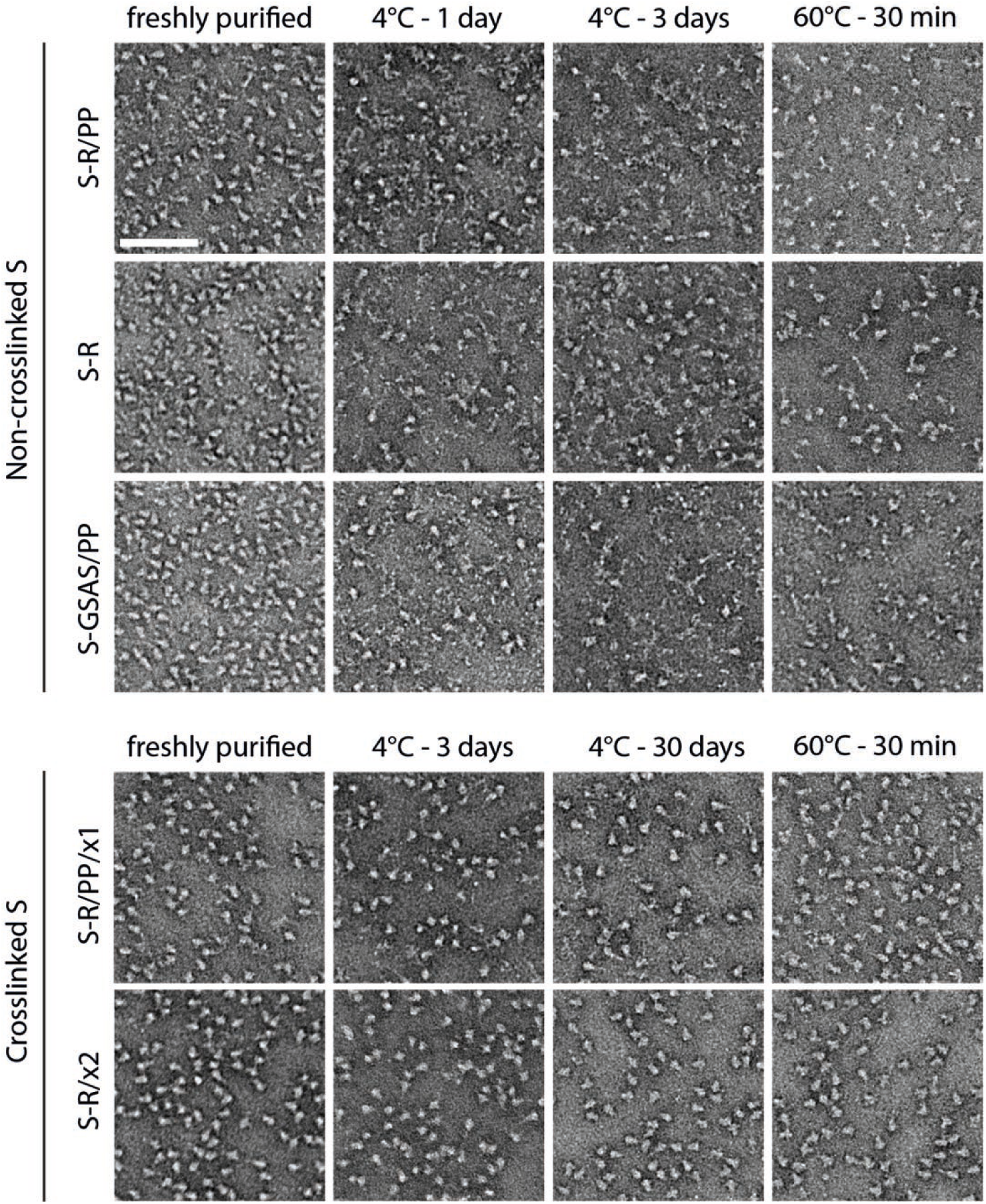
Oligomeric state and thermal stability of S protein trimers assessed by negative stain EM. Freshly purified proteins were imaged by negative stain EM (left column). Non-crosslinked S proteins show predominantly trimers with some smaller material, while S-R/x2 shows almost exclusively trimers. Upon prolonged incubation at 4°C, or upon heat treatment, non-crosslinked S proteins disintegrate, while crosslinked S proteins remain trimeric. Scale bar = 100 nm.

To assess the stability of the protein, we repeated the negative stain EM after prolonged storage at 4°C (**Figure 2, middle columns**). Non-crosslinked variants started to disintegrate into disordered structures after 1 day of storage, and disintegration is mostly complete after 3 days of storage. We suggest that the disintegration of spike trimers is facilitated by RBD opening. In contrast, the crosslinked variants could be stored at 4°C for at least 1 month without significant disintegration of trimers being observed by negative stain EM.

We further tested the stability of the crosslinked variants by subjecting them to incubation at 60°C for 30 mins before imaging by negative stain EM (**Figure 2, right column**). We found that non-crosslinked variants denatured upon exposure to 60°C while crosslinked mutants showed minimal changes, remaining in their trimeric state. These results indicate that stabilising S in a closed conformation by insertion of cross-links leads to a substantial increase in protein stability. S-R/x2 was arrested in the pre-fusion state by insertion of a single disulphide bond to prevent opening, without the insertion of any other stabilizing point mutations, suggesting that in the absence of crosslinking, S protein denaturation proceeds via transition into the open form.

### Structural features of crosslinked mutants

In order to understand the increased stability, we imaged S-R/PP/x1 and S-R/x2 by cryo-EM. For comparison we also performed cryo-EM on S-R/PP and S-R expressed and purified under the same conditions. Classification of the cryo-EM data show that S-R/PP/x1 and S-R/x2 exclusively form trimers with 3 closed RBDs, suggesting that the designed disulphides have successfully trapped the RBDs in the closed state (**Figure S2**). As expected, the disulphide bond is clearly resolved in the fully-crosslinked S-R/x2, and visible, but less-well resolved in the incompletely-crosslinked S-R/PP/x1 (**Figure 3a**). Both non-crosslinked constructs contain closed trimers, and trimers with one RBD in the open state (**Figure S2**). We observed that ∼ 20% of S-R/PP trimers are in the closed state and ∼ 80% have one open RBD (**Figure S2**). In contrast we found that ∼ 80% of S-R trimers are in the closed conformation and only ∼ 20% have one open RBD. We observed that in the closed S-R trimer, negatively charged residues D427 and D428 in the RBD are in proximity to positively charged K986 (**Figure 3a**), and may electrostatically stabilize the closed form. This electrostatic attraction is lost by substituting the basic K986 to neutral proline in S-R/PP. We suggest that the widely-used PP mutation favours the transition into the open state by abolishing electrostatic interactions between K986 and negatively charged residues in the RBD. The higher fraction of closed trimers present in S-R when compared to S-R/PP, provides an explanation for the higher efficiency of crosslinking seen in S-R/x2 when compared to S-R/PP/x1.

**Figure 3.**
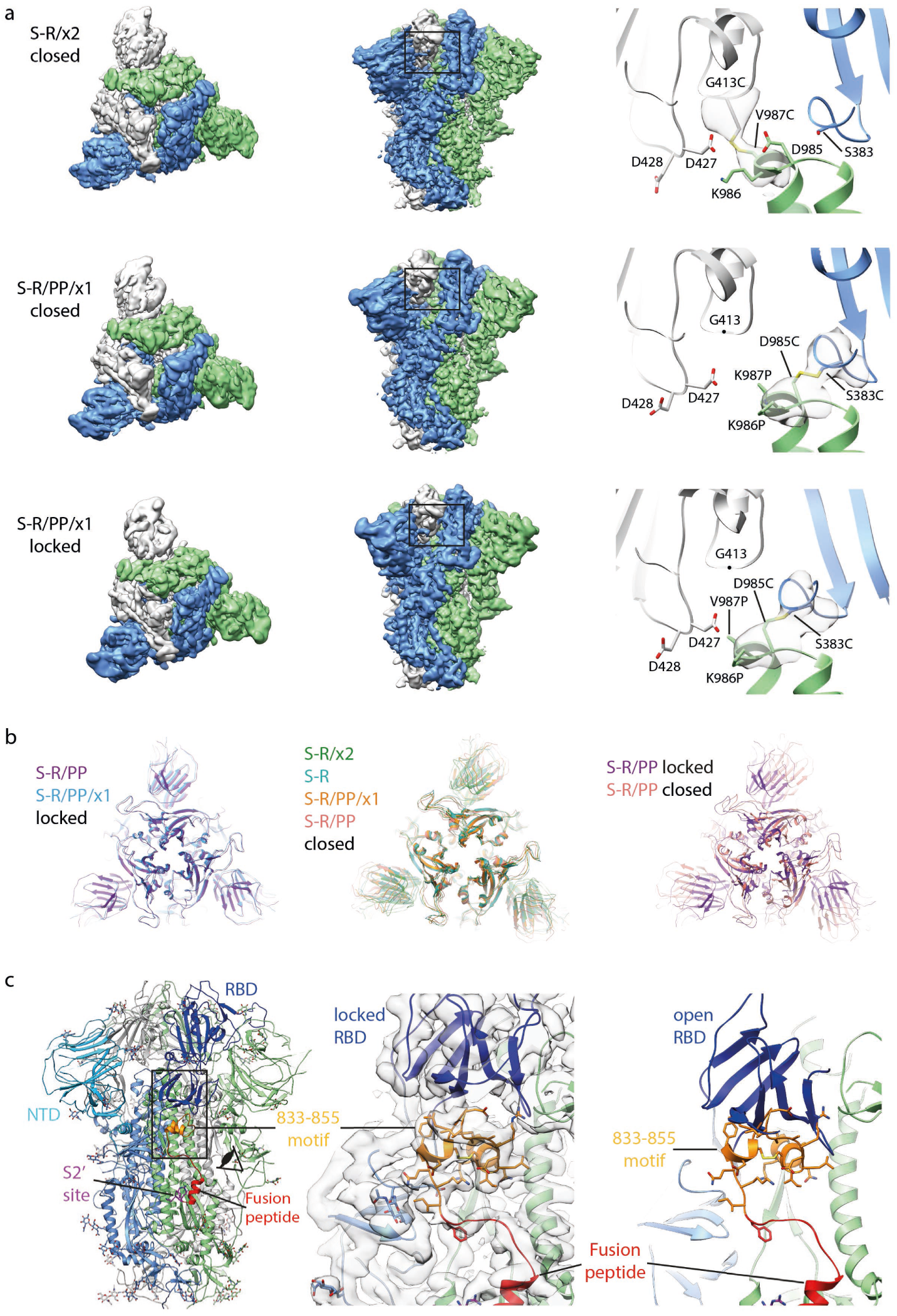
Characterisation of S-protein trimers by cryo-EM. **a)** Cryo-EM densities for S-R/x2 in the closed conformation, and for S-R/PP/x1 in both closed and locked conformations. Right hand panels show resolved density for the inserted disulphide bonds at the expected positions. **b)** Comparison of the RBD and NTD conformations in S trimers of locked and closed conformations as viewed from the top of the trimer at the three-fold axis after alignment based on the central S2 helices. Between the locked and closed conformations the S1 domains show a twist motion (right panel and compare left and middle panel). In the locked conformation, the RBD moves closer to the 3-fold axis. The position of the NTD is variable. **c)** Folding of 833-855 into a helix-turn-helix motif locks the RBD in a locked conformation. Left panel, locations of RBD, 833-855 motif (orange), fusion peptide and S2’ site are coloured and indicated. Middle panel, detailed density for the 833-855 motif is viewed from near the fusion peptide towards the RBD (indicated by the eye icon in the left panel). The density shows well resolved density that was disordered in previous SARS-CoV-1 and SARS-CoV-2 S structures. Right panel, the position of the RBD in the open conformation is aligned onto the locked monomer. In the open conformation the RBD would clash into the folded 833-855 motif (orange).

Further sub-classification of the closed conformations revealed that S-R and S-R/x2 adopt a single closed conformation similar to previously reported closed SARS-CoV-2 S structures, with some flexibility in the position of the NTD (residue 1-305) indicated by the lower resolution of this region in the EM density (**Figure S3a**). In contrast, S-R/PP and S-R/PP/x1 variants exist in two distinct closed states – while the majority of trimers are in the previously described closed conformation, some of the closed trimers are in a more tightly closed form (henceforth called locked conformation) in which the RBDs have twisted to move approximately 2 Å closer to the three-fold symmetry axis at the tip of the spike (**Figure 3b**).

The region N-terminal to the fusion peptide and S2’ site, residues 833-855, is disordered in the closed conformation (**Figure 3c**) and in all SARS-CoV-1 and SARS-CoV-2 S protein structures published to date. In the locked conformation it has folded into two helices connected by a loop (**Figure 3c**). Structural comparison to an open monomer suggests folding of this motif would prevent the motion of the RBD during opening, effectively locking the RBD in a closed conformation (**Figure 3c**). Consistent with this hypothesis, a folded motif is present at this position in the structures of S from murine hepatitis virus (MHV) ^22^, a betacoronavirus, and from alpha (NL63-CoV^23^), gamma (IBV, infectious bronchitis virus ^24^), and delta (porcine deltacoronavirus ^25^) coronaviruses – for these S proteins no open conformation was observed (**Figure S4a**). The NTD and RBD are better resolved in the cryo-EM density in the locked conformation than in the closed conformation (**Figure S3a**), suggesting they are less mobile in this conformation. A SARS-CoV-2 variant which has now become the prevalent form globally contains a D614G mutation ^26^ which would abolish a salt bridge between D614 and K854 within the folded 833-855 motif (**Figure S4b**). This change could potentially destabilise the locked conformation to promote RBD opening and potentially increase virus receptor binding and membrane fusion activities. We only observed the locked conformation in variants stabilized with PP, but a recent report of full-length, wild-type S trimers purified in detergent micelles ^11^ also described folding of the 833-855 region. This observation suggests that folded 833-855 is a relevant state that is selectively stabilised by the PP mutation. We suggest that the locked conformation represents an early intermediate state before RBD can open to bind receptor.

### Crosslinked proteins in diagnostic assays

We next compared the performance of S-R/x2 and non-crosslinked constructs in diagnostic assays. First, we performed ELISA titrations against S-R/x2 for sera from forty-eight individuals who are seropositive against S-R/PP. A positive response against S-R/x2 was seen for forty-two of these sera. There was a clear correlation between the strength of response against S-R/PP and S-R/x2 (**Figure 4a**), with the response against S-R/x2 reduced by approximately 3-fold (**Figure 4b**). This reduction presumably reflects the reduced exposure of RBD epitopes in the closed conformation. Second, we performed Luminex-based assessment of the binding of S-GSAS/PP, S-R/PP, S-R, S-R/x2 and isolated RBD to IgG in sera from seven other individuals who are seropositive against SARS-CoV-2 antigens (four patients, and three individuals with a history of mild or no symptoms). A strong response was seen for all five constructs against all seven sera (**Figure 4c**). S-GSAS/PP and S-R/PP showed strongest and almost identical responses, S-R showed a slightly lower and S-R/x2 a reduced response, in the latter two cases presumably due to the reduced exposure of epitopes within the ACE2-binding region of the RBD in the closed spike conformation. The response to S-R/x2 is considerably stronger than that to RBD, suggesting that the closed S trimer displays considerably more epitopes than RBD alone.

**Figure 4.**
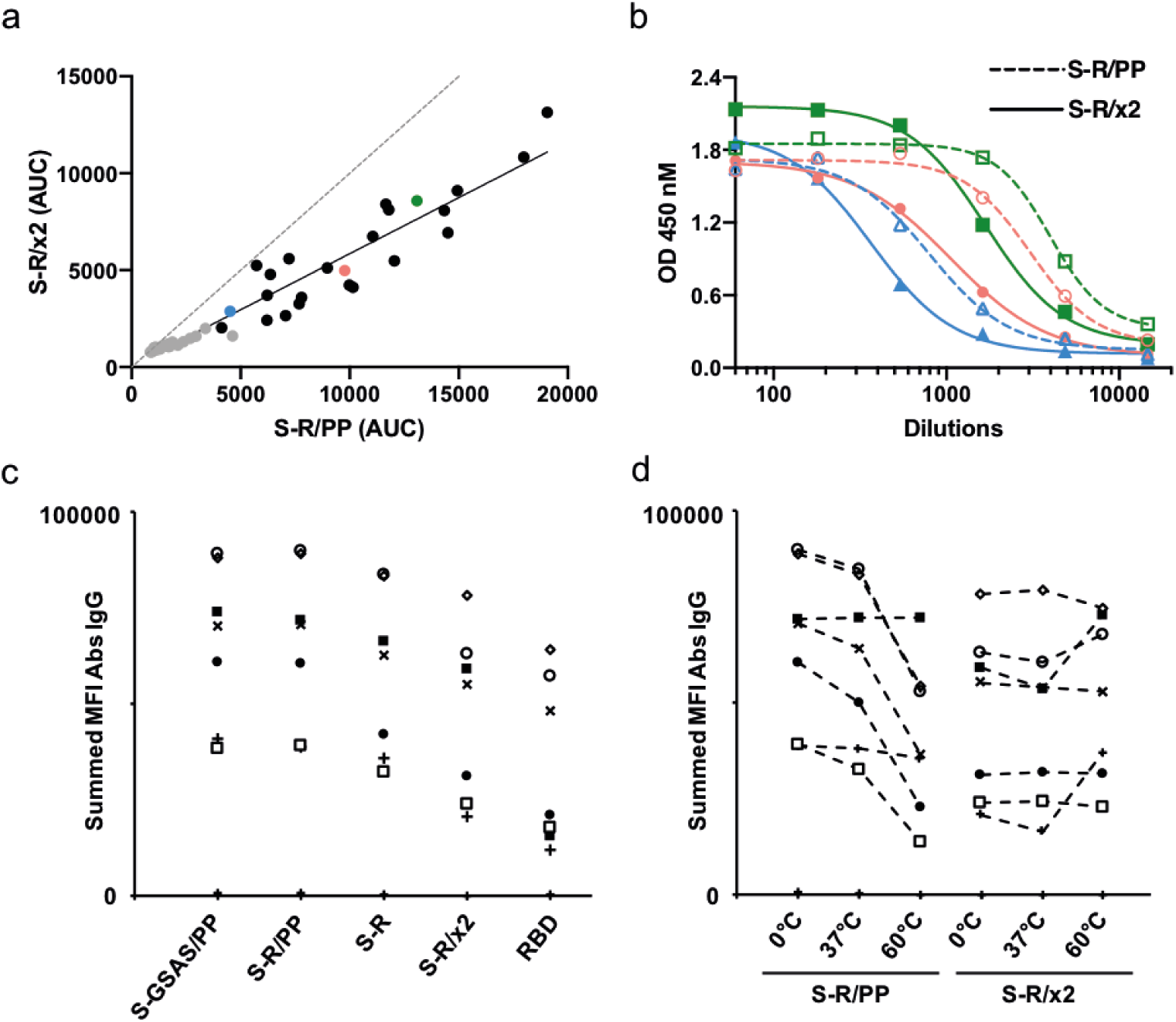
Behaviour of crosslinked proteins in diagnostic assays. **a-b)** Comparison of immune responses to open and closed conformations of the SARS-CoV-2 spike protein. Serum samples from PCR positive patients were screened for reactivity against S-R/PP and any samples scored as positive (n=48) were subsequently selected for end-point titrations against both for S-R/PP and S-R /x2. **a)**. The area under the curve (AUC) was calculated for each seropositive sample using end point titrations. Paired AUC values for reactivity against S-R/PP and S-R /x2 were plotted for each positive sample. Pearson’s correlation analysis was calculated as r=0.9599 (95% CI 0.929 – 0.977) R^2^ of 0.921. The dashed line represents the theoretical line of equality, whereas the solid line represents the best fit using linear regression. Where possible, samples were used for subsequent EC50 determination (**Figure S5**). Data points in grey are those where an EC50 for both S-R/PP and S-R /x2 could not be determined as one or both did not reach saturation. Coloured data points highlight three representative samples shown in panel b. **b)** End point titration curves against S-R/PP and S-R /x2. In each case the dashed line represents reactivity against the open conformation S-R/PP and the solid line represent the reactivity against the closed conformation S-R /x2. **c-d)** Summed mean fluorescence intensity (MFI) in a multiplexed flow cytometry (Luminex) assay to measure binding of IgG in sera to S protein variants and RBD. Sera from seven individuals seropositive for SARS-CoV-2 are shown, as well as two negative control sera (points along the x-axis). In d), S proteins were incubated at the indicated temperatures prior to use in the assay.

We used the Luminex-based assay to assess the impact of S protein conformational state on interactions with antibodies in sera. We incubated S-R/PP and S-R/x2 at 37°C for 36 hours, or subjected them to 60°C for 30 minutes, prior to coupling to Luminex beads. We then compared their binding to the same seven sera used above (**Figure 4d**). In the case of S-R/PP we found that exposure to temperature leads to a reduction in IgG response, consistent with the loss of trimeric structure observed by negative stain EM (**Figure 2**), for five sera. In contrast, S-R/x2, which maintains its structure upon heat treatment, also maintains its antibody interactions with sera for these five sera. Two sera did not show a loss of binding upon heat treatment of S-R/PP, and showed increased binding against 60°C heat-treated S-R/x2. Notably these two sera also have the lowest binding against RBD. We cannot draw conclusions on the serological differences between the individuals based on this data but speculate that they may contain antibodies against peripheral linear epitopes which become increasingly exposed upon heat treatment.

The behaviour of S-R/x2 in diagnostic ELISA and Luminex assays therefore correlated well with its structural behaviour. S-R/x2 has a generally lower response than the proline-stabilized S-R/PP, consistent with the fact that it hides epitopes in the RBD. The S-R/x2 response is in most cases not altered by heat treatment, while the response of S-R/PP generally drops upon heat treatment, concomitant with structural denaturation. We believe that thermostabilization of S by insertion of x2 may have direct applications in production of more stable S derivatives for vaccine and diagnostic purposes.

### Potential applications of closed trimeric S constructs

The pattern of immune response against SARS-CoV-2 is not yet fully understood and varies between individuals. In this context an exclusively closed trimer with excellent stability like S-R/x2 has a number of potential applications. First, although we saw a correlation between response against S-R/PP and S-R/x2, there is variation in the relative responses (Figure S5) which likely reflect different antibody compositions in the sera. A recent preprint demonstrates that some monoclonals bind differently to an open trimer when compared to a closed trimer similar to our construct S-GSAS/PP/x1 ^20^. Exclusively closed trimers may therefore prove useful as a serological reagent to assay differential immune responses.

Second, while SARS-CoV-2 neutralising antibodies targeting alternative epitopes have been identified ^27^, the majority of neutralizing monoclonals identified to date act by blocking the interaction between the RBD and the receptor ACE2 ^28–30^. Antibodies that recognise the ACE2 binding site or cryptic epitopes in the open trimer can functionally mimic the receptor to trigger dissociation of S1 and S2 transition into the postfusion state ^8,18,31^. It is not clear how the resulting S-protein products might modulate a further immune response. In addition, binding of IgG to the MERS S receptor binding site and concomitant Fc-Fc receptor engagement promotes activation and internalization of the virus causing antibody enhancement of infection ^32^. We find that S-R/PP or related, published PP-containing constructs, exhibit a higher fraction of open states than the closer-to-wild type S-R. Exclusively closed trimers such as S-R/x2, if used as an antigen in vaccination or immunization, can be expected to induce an immune response that largely excludes receptor binding site targeting antibodies and thereby will favour alternative antibodies which do not compete or functionally mimic ACE2 and which may have the potential to lock S protein in a prefusion state to prevent infection. The stabilization of alternate or transient states in order to induce alternative immune responses has been a valuable approach in the elicitation of broadly neutralizing antibodies against HIV-1 Env, for example ^33^.

We are optimistic that S-R/x2, or derivatives thereof, will be applicable as diagnostic reagents, as immunogens, and as tools to generate or characterize novel monoclonal antibodies for immunotherapy.

## Methods

### Expression Constructs

DNA constructs were designed to transiently express ectodomains of SARS-CoV-2 S protein for secretion into cell culture media. The DNA sequence for the ectodomain (residues 14-1211) of S from 2019-nCoV BetaCoV/Wuhan/WIV04/2019 (GISAID accession no. EPI_ISL_402124) ^1^, covering the region from the end of signal peptide sequence (starting with QCVN) to the beginning of the transmembrane domain (ending with QYIK), was synthesised and codon optimised for HEK293 human cell expression. We added a Kozak sequence as well as sequences encoding an exogenous N-terminal signal peptide MGILPSPGMPALLSLVSLLSVLLMGCVAETGT derived from µ-phosphatase and a C-terminal extension GSGR*ENLYFQ*GGG<u>GSGYIPEAPRDGQAYVRKDGEWVLLSTFL</u>G**HHHHHH**. The C-terminal extension contains a TEV site (italic), a T4 trimerization foldon (underlined) and a Hexa-histag (bold). The extended sequence was cloned into a pCDNA3.1 vector. Modification of the coding sequence of the multibasic S1/S2 cleavage site PRRAR to PGSAS or to a single R was carried out by PCR mutagenesis using Q5 polymerase (New England Biolabs). Introduction of cysteine crosslinks and modification of residues 986 and 987 were carried out using Q5 polymerase PCR with primers containing desired substitutions, followed by In-Fusion HD (Takara Bio) assembly. Cysteine positions were designed based on visual inspection of published structures, and using Disulfide by Design 2.0 ^34^.

### Protein Production and Purification

Expi293 cells (Thermo Fisher) cultured in Expi293 media (Thermo Fisher) were maintained at 37°C. Cells were diluted to a density of 2-3×10^6^ cells/ml before transfection. For protein production, 500 µg of DNA was mixed with 1350 µg of polyethylenimine in 10 ml Expi293 media, incubated for 10 min, then added to 500 ml of cells. Transfected cells were cultured at 33°C. Except where otherwise indicated, for non-cystine crosslinked constructs, transfected cell culture supernatant was harvested twice at day 3 and day 6. For cysteine-crosslinked mutants, supernatant was harvested once at day 6.

To purify S protein ectodomains, cells were removed from culture media by centrifugation and 10X phosphate buffered saline (PBS) was added to 1X concentration. Cleared media was supplemented with 2.5 mM imidazole, 300 mM NaCl and 0.5 mM phenylmethylsulfonyl fluoride and recirculated onto a 5 ml Talon Cobalt column 2.5 times. After sample application, the column was washed by 50 ml buffer A (25 mM Phosphate pH 8.0, 300 mM NaCl, 5 mM Imidazole), and protein was eluted with a 100 ml linear gradient to 100% buffer B (25 mM Phosphate pH 8.0, 300 mM NaCl, 500 mM Imidazole). Fractions containing S protein were pooled, proteins were concentrated and buffer exchanged into PBS in a 100 kDa MWCO spin concentrator. Purification from 500 ml cells typically yielded 150-200 µg of crosslinked protein, ∼700 µg of S-R/PP and ∼300 µg of S-R. Except where otherwise indicated, proteins were flash frozen in liquid nitrogen and stored at −80°C.

### Western Blot

30 µl cleared cell culture supernatant ∼80 hours post-transfection was heat denatured by addition of 4X NuPAGE™ LDS Sample Buffer (Thermo Fisher) in non-reducing and reducing (in the presence of 250 mM beta-mercaptoethanol) conditions at 90°C for 5 min. Protein were separated by electrophoresis in NuPAGE™ 4% to 12%, Bis-Tris gels (Thermo Fisher). Protein bands were transferred onto Trans-Blot Turbo Midi 0.2 µm PVDF membranes (BioRad) in a Trans-Blot Turbo Transfer System (BioRad). Membranes were blocked with 5% skim milk in PBS supplemented with 0.1% Tween 20. Presence of His-tagged protein was detected by chemical luminescence using His-probe (H-3) HRP (sc-8036 HRP, Santa Cruz Biotechnology) monoclonal antibody horseradish peroxidase (HRP) conjugate and Pierce™ ECL Western Blotting Substrate (Thermo Fisher).

### Negative staining EM

Proteins were diluted to ∼0.05 mg/ml in water. CF200-Cu carbon film grids (EMS) were glow discharged for 45 seconds at 25 mA in air, 3 µl of diluted protein was applied and absorbed for 1 min. The grids were side-blotted and washed once in water and stained twice with Nano-W stain (Nanoprobes) with blotting in between. The grids were air dried and imaged using a Tecnai T12 microscope operated at 120 kV.

### Cryo-EM

Grid preparation and image collection were performed similarly for all purified native, mutated and cross-linked SARS2 S proteins. C-Flat 2/2 3C grids (Protochips) were glow-discharged for 45 seconds at 25 mA. 3 µl of freshly purified protein at ∼0.2-0.6 mg/ml supplemented with 0.01% octyl-glucoside (OG) was applied to the grids, which were plunge-frozen in liquid ethane using a Vitrobot (Thermo Fisher Scientific). Double loading with side-blotting between loading was performed when the concentration of purified protein was below 0.3 mg/ml (S-R/x2). Grids were stored in liquid nitrogen and loaded into a Titan Krios electron microscope (Thermo Fisher Scientific) operated at 300kV, equipped with a Gatan K3 BioQuantum direct electron detector with the slit retracted. Movies with 48 frames and an accumulated dose of 50 electrons/Å^2^ were acquired in counting mode using SerialEM-3.8.0 ^35^ at the magnification of 81,000 X, corresponding to a calibrated pixel size of 1.098 Å/pixel. Detailed data acquisition parameters are summarized in Extended Data Table 1.

### CryoEM image processing

RELION-3.1’s Scheduler (to be described elsewhere) was used for fully automated real-time processing up to initial 3D classification while micrographs were being collected on the microscope. In a first Schedule called *preprocess*, movies were motion-corrected and dose-weighted using RELION’s implementation of the MotionCor2 algorithm ^36^; the contrast transfer function (CTF) was estimated by CTFFIND-4.1.13 ^37^; and particles were automatically picked by template matching using a structure of the S protein filtered to 20 Å resolution as a 3D reference. In parallel, a second Schedule called *class3d* used the particles generated by the *preprocess* Schedule for 3D classification in batches of 500,000 particles without imposing symmetry to separate SARS-Cov2 S protein particles from contaminating features.

The remainder of the processing was performed interactively. Particles belonging to 3D classes corresponding to S-protein particles were subjected to one round of 2D classification to remove any remaining contaminating features. Subsequently, a second round of 3D classification was used to assess the ratio of open and closed states and standard 3D auto-refinement was performed on pooled classes corresponding to the different states (Fig. S2). Suitable closed and locked classes were subjected to Bayesian polishing to correct for per-particle beam-induced motions, followed by CTF refinement for per-particle defocus, per-micrograph astigmatism, beam-tilt and optical aberration correction ^38,39^ and a final round of 3D auto-refinement.

Map resolutions were estimated at the 0.143 criterion of the phase-randomization-corrected FSC curve calculated between two independently refined half-maps multiplied by a soft-edged solvent mask. Final reconstructions were sharpened and locally filtered in RELION *post-processing* (Fig. S3). The estimated B-factors of each map are listed in Extended Data Table 1.

### Model building and refinement

The SARS-CoV-2 S protein ectodomain structure (PDBID: 6VXX ^19^) was used as a starting model for all closed conformations. Models were built and adjusted manually in Coot ^40^. Steric clash and sidechain rotamer conformations were improved using the Namdinator web server ^41^. After further manual adjustment, the structures were refined in PHENIX-1.18.261 ^42^ to good geometry and statistics are given in Extended Data Table 1. The unmasked model-to-map FSC was calculated in PHENIX for the refined model against the full reconstruction to validate the confidence of the atomic model.

### ELISA assays

The ELISA assay was performed in a two-step process whereby a positivity screen was then followed by an endpoint titer determination following previously established protocols ^43^. Briefly, 96-well EIA/RIA plates (Corning, Sigma) were coated with PBS or 0.1μg per well of either S-R/PP or S-R/x2 at 4°C overnight. On the following day, the coating solution was removed and wells were blocked with 3% skimmed milk prepared in PBS with 0.1% Tween 20 (PBST) at room temperature for 1 h. Serum samples previously inactivated by heating at 56°C for 1 h were diluted to 1:60 or serially diluted by 3-fold, six times in 1% skimmed milk in PBST. The blocking solution was removed and the diluted sera was added to the plates and incubated for 2 h at room temperature. Following the incubation, the diluted sera were removed and the plates were washed three times with PBST. Goat anti-human IgG secondary antibody-Peroxidase (Fc-specific, Sigma) prepared at 1:3000 in PBST was then added and plates were incubated for 1 h at room temperature. The plates were then washed three times with PBST. The ELISA was developed using 3,5,3′,5′-tetramethylbenzidine (TMB, Thermo Scientific) solution and the reaction was stopped after 10 min incubation using 0.16M Sulfuric acid. The optical density at 450 nm (OD450) was measured using a Spectramax i3 plate reader.

The absorbance values for each sample were determined after subtracting the OD values from uncoated wells. All data analyses were performed using Prism 8 version 8.4.2 (GraphPad). To investigate the impact of conformation on reactivity responses, area under the curve (AUC) values for all positive samples were calculated and Pearson’s correlation analysis comparing responses to S-R/PP and S-R/x2 was calculated. In addition, the dilution of sero-positive samples required to give a 50% maximum signal (EC_^50^_) against different spike protein constructs was determined using non-linear regression analysis in Prism 8 version 8.4.2.

Serum samples for the ELISA assays were obtained from patients attending Addenbrooke’s Hospital with a suspected or confirmed diagnosis of COVID19 together with Health Care Workers identified through the staff screening as PCR positive for SARS-CoV-2. Ethical approval was obtained from the East of England – Cambridge Central Research Ethics Committee (REC ref 17/EE/0025). All participants provided informed consent.

### Multiplex particle-based flow cytometry (Luminex)

S-GSAS/PP, S-R/PP, S-R, S-R/x2, RBD were covalently coupled to distinctive carboxylated bead sets (Luminex; Netherlands) to form a 5-plex assay. The RBD protein used in this assay includes residues 319-541 with addition of the first 14 residues of S to the N-terminus and a 6XHis tag to the C-terminus ^44^ It was expressed and purified from HEK293 cells.

Beads were first activated with 1-ethyl-3-[3-dimethylaminopropyl]carbodiimide hydrochloride (Thermo Fisher Scientific) in the presence of N-hydroxysuccinimide (Thermo Fisher Scientific), according to the manufacturer’s instructions, to form amine-reactive intermediates. The activated beadsets were incubated with the corresponding proteins at a concentration of 50 μg/ml in the reaction mixture for 3 h at room temperature on a rotator. Beads were washed and stored in a blocking buffer (10 mM PBS, 1% BSA, 0.05% NaN3).

The S-variant and RBD coupled bead sets were incubated with SARS-CoV-2 patient sera at 3 dilutions (1/100; 1/1000 and 1/10000) for 1 h in 96-well filter plates (MultiScreenHTS; Millipore) at room temperature in the dark on a horizontal shaker. Fluids were aspirated with a vacuum manifold and beads were washed three times with 10 mM PBS/0.05% Tween 20. Beads were incubated for 30 min with a PE-labeled anti–human IgG-Fc antibody (Leinco/Biotrend), washed as described above, and resuspended in 100 μl PBS/Tween. They were then analyzed on a Luminex analyzer (Luminex / R&D Systems) using Exponent Software V31. Specific binding was reported as mean fluorescence intensities (MFI). MFI values at each of the three dilutions were summed to generate the points shown in Figure 4e,f as representative values for comparison purposes.

Patient sera for Luminex assays were collected following informed consent, aliquoted and held at −70°C prior to use. Samples were taken from patients with PCR-confirmed COVID19 looked after at Royal Papworth Hospital NHS Foundation Trust (RPH), and RPH healthy staff who presented with either no history or a mild history of COVID19 symptoms. Samples screened positive for SARS-CoV-2 N and S binding antibodies by Luminex assay as described were selected for further investigation. Ethics Approval: IRAS Project ID: 96194 REC: 12/WA/0148.

## Acknowledgements

We thank the staff of the MRC-LMB for generous support during the COVID-19 pandemic lockdown. We thank the staff of the MRC-LMB EM Facility, in particular Grigory Sharov and Giuseppe Cannone, for supporting EM; Jake Grimmett and Toby Darling for supporting scientific computing; and Patricia Edwards for supporting cell culture. We thank Takanori Nakane for advice on image processing. We thank Donna Mallery and David Paul for performing exploratory experiments. We thank the Royal Papworth Hospital Clinical Staff and R&D team in supporting patient and staff recruitment, J. Gronlund for support in sample collection and Royal Papworth Hospital healthy staff donors for their participation in this research. This study was supported by funding from the European Research Council (ERC) under the European Union’s Horizon 2020 research and innovation programme (ERC-CoG-648432 MEMBRANEFUSION to JAGB), the Medical Research Council as part of United Kingdom Research and Innovation (MC_UP_A025_1011 to APC; MC_UP_A025_1013 to SHWS; MC_U105181010 to LCJ; MC_UP_1201/16 to JAGB), 100 Top Talents Program of Sun Yat-sen University (to YZ), a Wellcome Trust Senior Fellowship (207498/Z/17/Z to IG).

## Author contributions

XX conceived the project. XX and JAGB planned the study. XX and YZ designed and synthesized the initial expression construct. XX, APC and JAGB designed crosslinked and mutated constructs which were made by APC. Protein purification was established by XX and performed by XX and KAC. XX performed negative stain and prepared EM grids. KQ and ZK performed cryo-EM, SHWS assisted with RELION scheduler, KQ processed EM data which XX, KQ and JAGB interpreted. Recruitment of donors, management of sera collection and sera processing were performed by LB, SB, CNCBRC and HEB. ELISA assays were performed by MH and IG, data analysis by MH, IG and JAGB. Luminex assays were designed by RD, performed by SE, data analysis by RD and JAGB. GLG and JAN provided critical reagents. LCJ contributed to design and interpretation. AJC, SHWS, YZ, LJC, IG and JAGB obtained funding. JAGB supervised and managed the project. XX, KQ, MH and JAGB made figures. XX, KQ and JAGB wrote the paper with input from all authors.

## The CITIID-NIHR COVID-19 BioResource Collaboration

**Principal Investigators:** John Bradley, Paul A. Lyons, Kenneth G.C. Smith, Mark Toshner. **CRF and Volunteer Research Nurses:** Anne Elmer, Carla Ribeiro, Jenny Kourampa, Sherly Jose, Jane Kennet, Jane Rowlands, Anne Meadows, Criona O’Brien, Rebecca Rastall, Cherry Crucusio, Sarah Hewitt, Jane Price, Jo Calder, Laura Canna, Ashlea Bucke, Hugo Tordesillas, Julie Harris, Valentina Ruffolo, Jason Domingo, Barbara Graves, Helen Butcher, Daniela Caputo and Emma Le Gresley. **Sample Logistics:** Benjamin J Dunmore, Jennifer Martin, Ekaterina Legchenko, Carmen Treacy, Christopher Huang, Jennifer Wood, Rachel Sutcliffe, Josh Hodgson, Joy Shih, Stefan Graf, Zhen Tong, Federica Mescia, Tobias Tilly, Ciara O’Donnell, Kelvin Hunter, Linda Pointon, Nicole Pond, Marta Wylot, Emma Jones, Stuart Fawke and Ben Bullman. **Sample Processing:** Laura Bergamaschi, Lori Turner, Isobel Jarvis, Ommar Omarjee, Aloka De Sa, Joe Marsden, Ariana Betancourt, Marianne Perera, Maddie Epping, Nathan Richoz, Georgie Bower, Rahul Sharma, Francesca Nice, Oisin Huhn and Stuart Fawke. **NIHR BioResource:** Hannah Stark, Neil Walker, Kathy Stirrups, Nigel Ovington, Eleanor Dewhust, Emily Li and Sofia Papadia.

---

**Figure S1.**
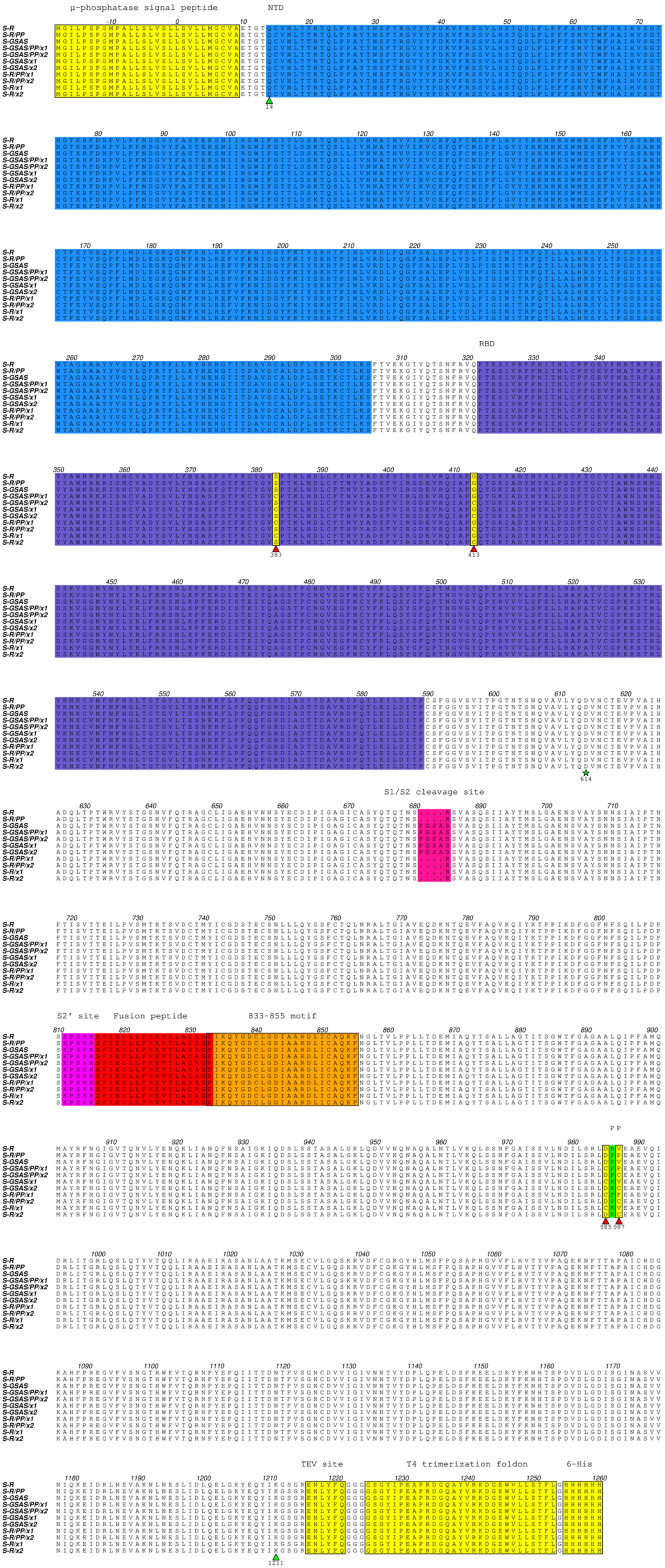
Amino acid sequences of constructs used in this study. The beginning (residue 14) and the end (1211) of native S protein ectodomain cloned in this construct are indicated with green triangles. Amino acid numbers were offset such that they correspond to the native sequence numbering. Extensions to the protein including the µ-phosphatase signal peptide, TEV site, T4 trimerization foldon and 6-His tag are coloured yellow and indicated. Structural elements including NTD, RBD, S1/S2 cleavage site, S2’ site, fusion peptide, and the 833-855 motif identified in this study are coloured and indicated. Sites which are modified in different constructs, including 383, 413, 985, 987, the S1/S2 cleavage site, and the double proline site (PP) are coloured and indicated. Positions at which cysteines were inserted are marked with red triangles. The site of D614G variation which is prevalent in the current circulating SARS-CoV-2 strain is indicated with a star.

**Figure S2.**
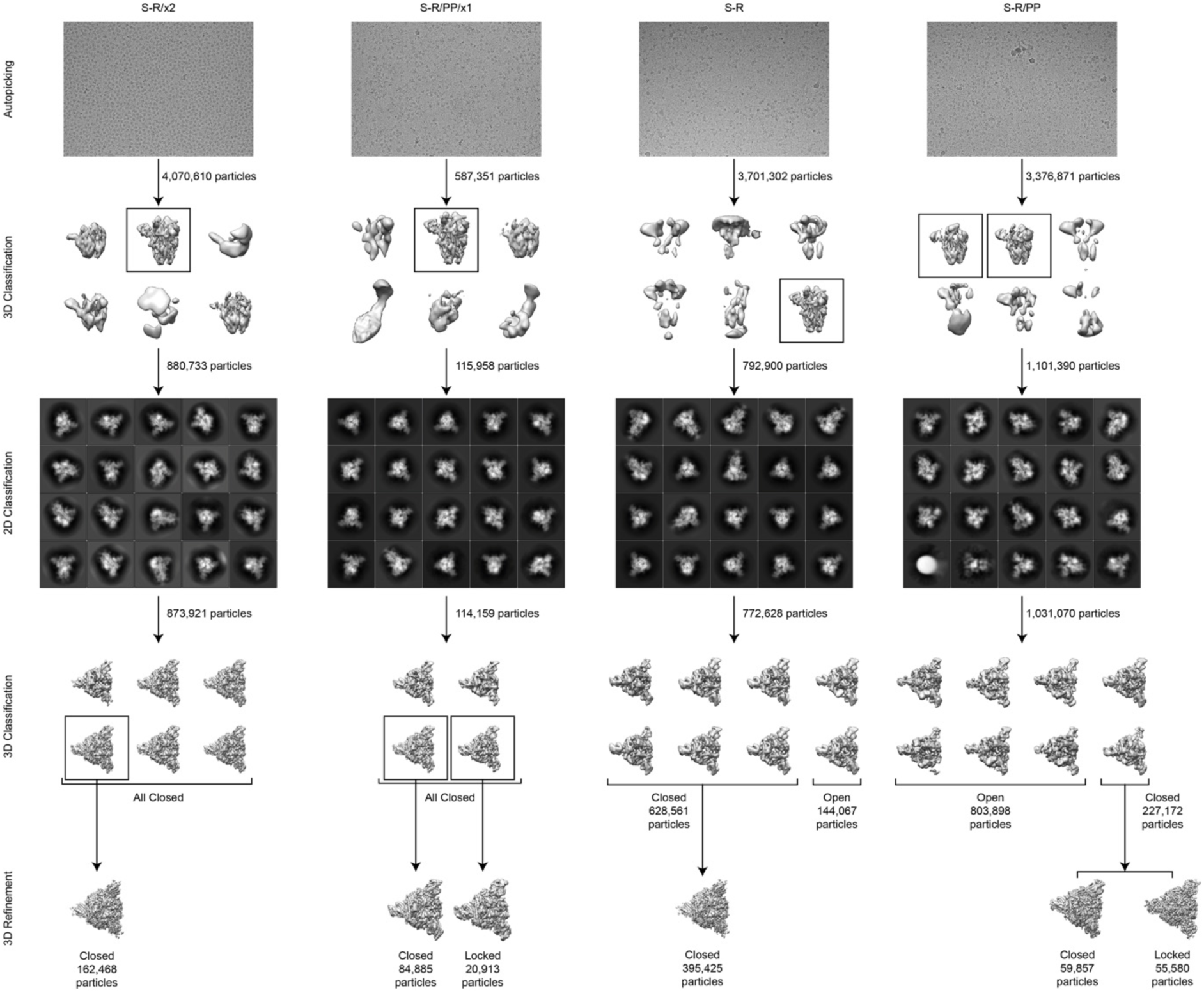
Pipeline used for picking and classification of cryo-EM data for S-R/x2, S-R/PP/x1, S-R, and S-R/PP. After automated picking, 3D and 2D classification steps were used to remove contaminating objects. 3D classification was then used to sort the data into open, closed and locked conformations.

**Figure S3.**
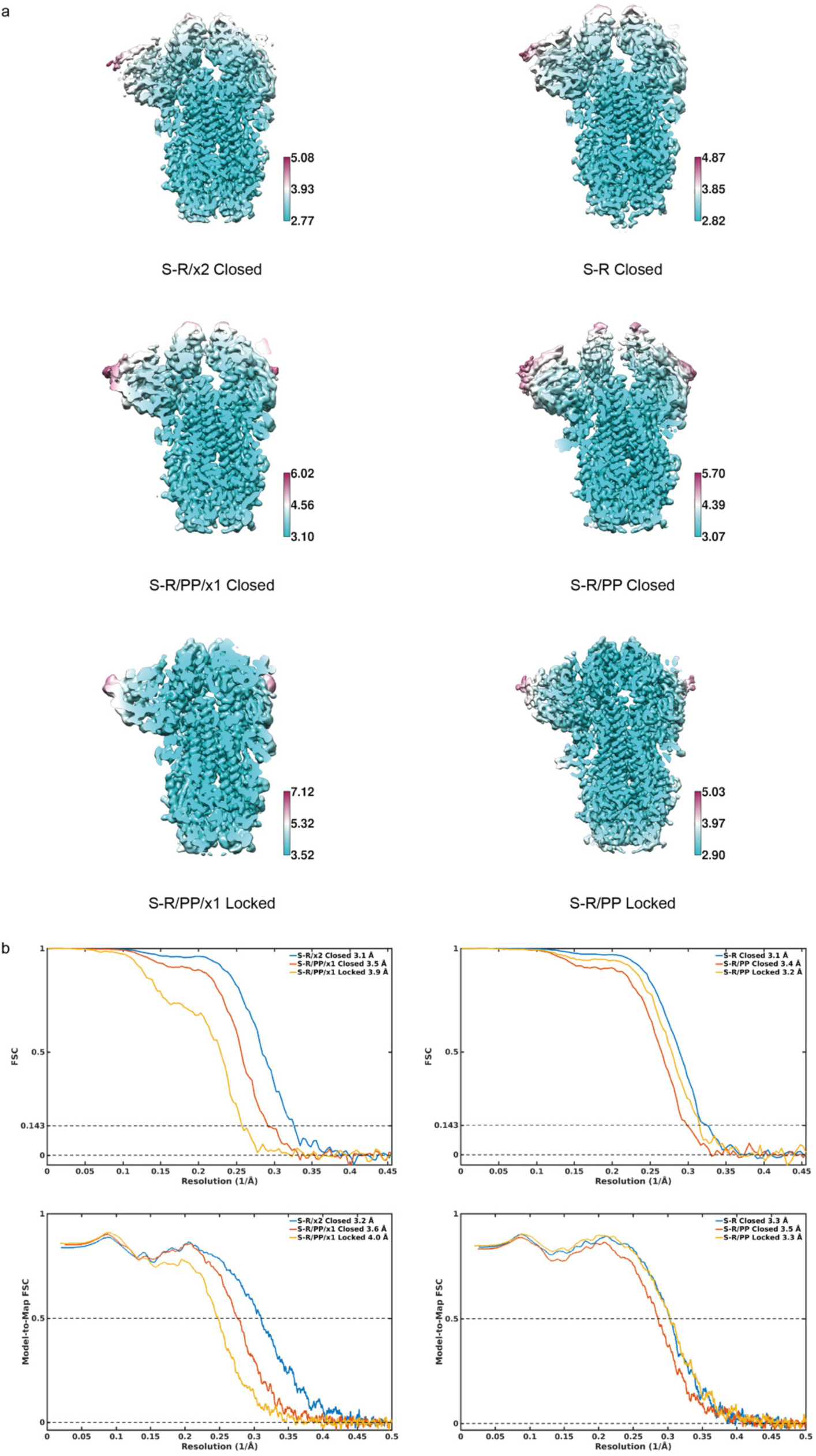
Resolution assessment of cryo-EM structures. **a)** Local resolution maps for all closed structures calculated using RELION. **b)** Global resolution assessment by Fourier shell correlation at the 0.143 criterion, and correlations of model vs map by Fourier shell correlation at the 0.5 criterion.

**Figure S4.**
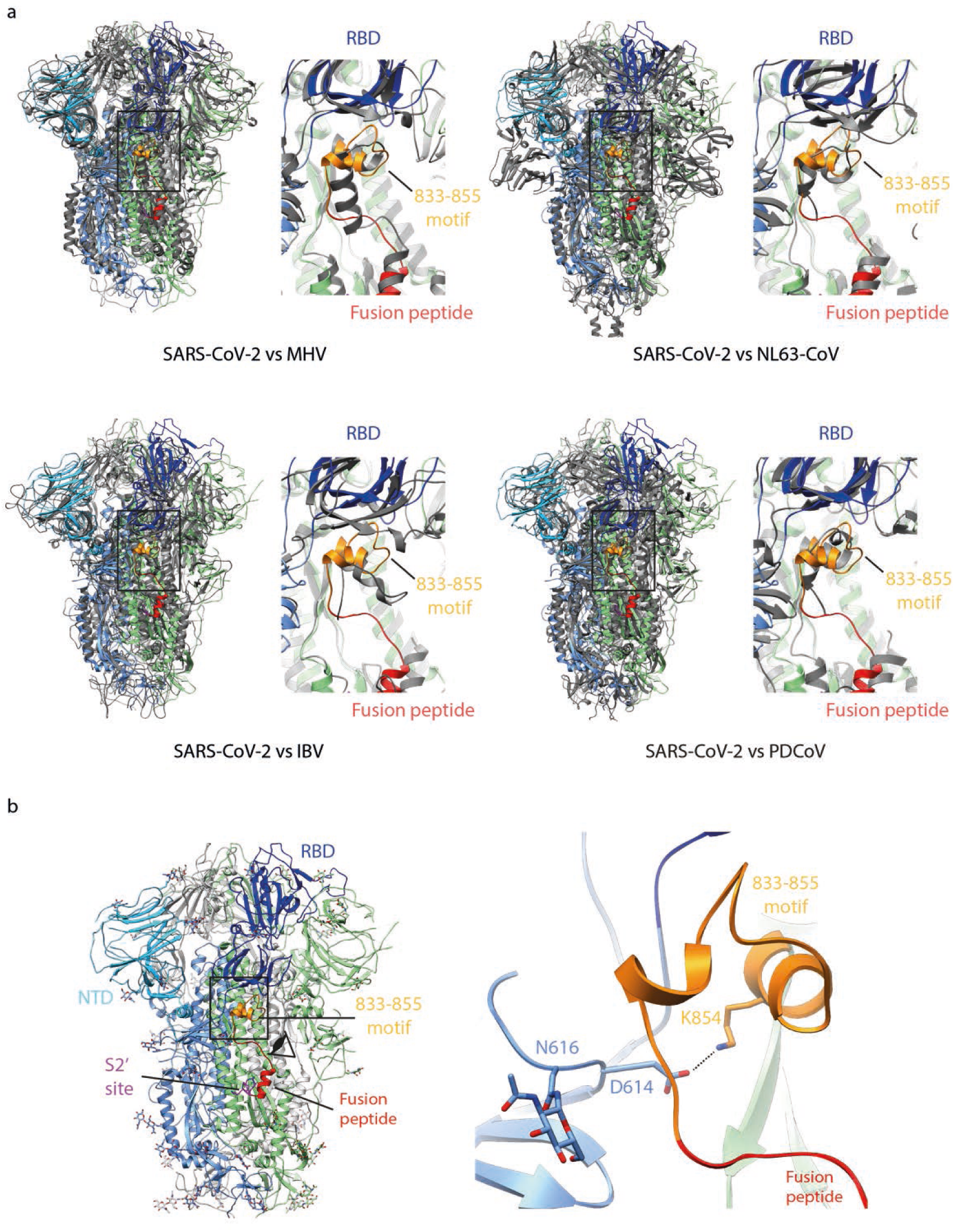
Structure of the 833-855 motif. **a)** Structural comparison with other closed coronaviruses spikes at the 833-855 motif. SARS-CoV-2 S is compared to spike proteins of murine hepatitis virus (MHV, a betacoronavirus, PDBID: 3JCL), NL63 coronavirus (NL63-CoV, an alphacoronavirus, PDBID: 5SZS), infectious bronchitis virus (IBV, a gammacoronavirus, PDBID: 6CV0) and porcine deltacoronavirus (PDCoV, a deltacoronavirus, PDBID: 6BFU). S proteins are structurally aligned based on S2. S protein trimers from all 4 genera of coronaviruses show presence of folded motifs equivalent to the 833-855 motif identified in SARS-CoV-2 S, suggesting a universal mechanism of locking RBD in closed state in coronavirus spikes. **b)** Interaction between residue D614 and the 833855 motif by formation of a 2.S Å salt bridge between D614 and K854. Surrounding structural elements, including fusion peptide and glycans at N616 are shown and indicated.

**Figure S5:**
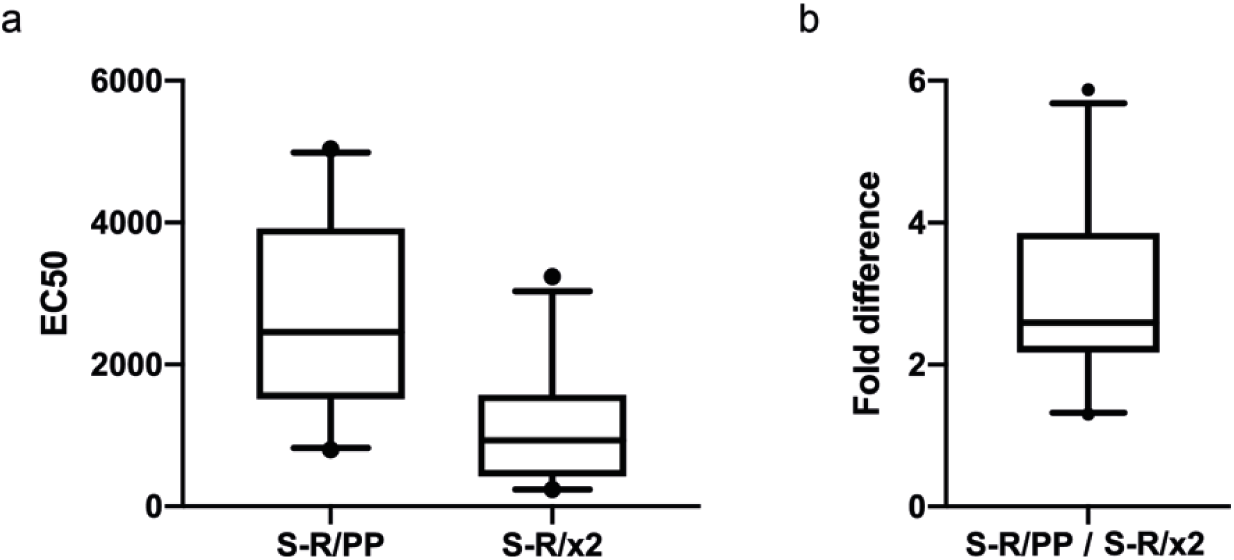
Comparison of immune responses to open and closed conformations of the SARS CoV-2 spike protein. **a)** The dilution of sero-positive samples required to give a 50% maximum signal (EC_50_) against S-R/PP and S-R/x2 was determined using non-linear regression analysis (n=24). The mean and the S5% confidence intervals are shown. The fold difference in reactivity against S-R/PP and S-R/x2 is shown in panel **b**

**Table S1.**
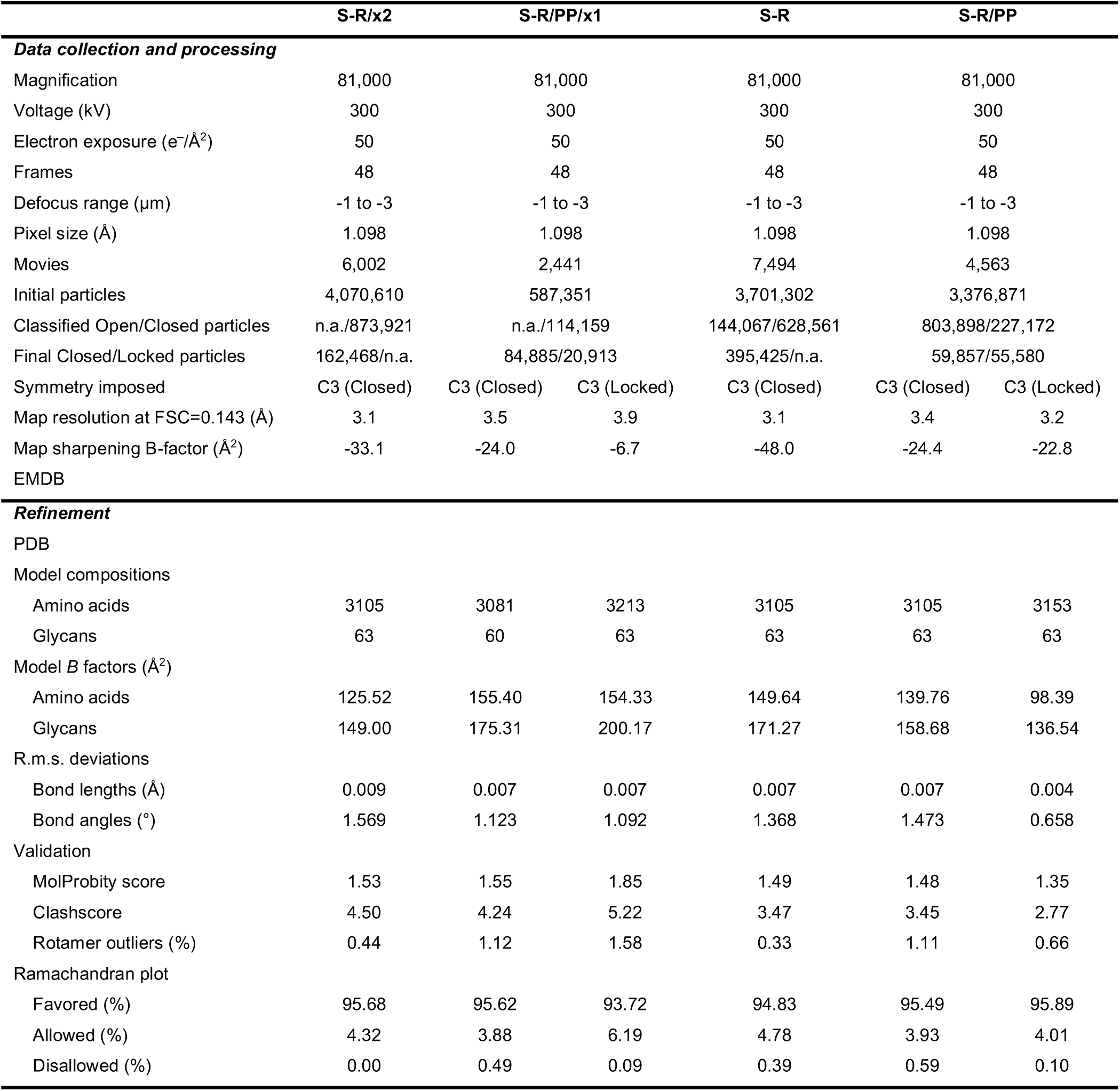
Cryo-EM data collection and refinement statistics.

|  | S-R/x2 | S-R/PP/x1 |  | S-R | S-R/PP |  |
| --- | --- | --- | --- | --- | --- | --- |
| **Data collection and processing** |  |  |  |  |  |  |
| Magnification | 81,000 | 81,000 |  | 81,000 | 81,000 |  |
| Voltage (kV) | 300 | 300 |  | 300 | 300 |  |
| Electron exposure (e<sup>-</sup>/Å<sup>2</sup>) | 50 | 50 |  | 50 | 50 |  |
| Frames | 48 | 48 |  | 48 | 48 |  |
| Defocus range (μm) | -1 to -3 | -1 to -3 |  | -1 to -3 | -1 to -3 |  |
| Pixel size (Å) | 1.098 | 1.098 |  | 1.098 | 1.098 |  |
| Movies | 6,002 | 2,441 |  | 7,494 | 4,563 |  |
| Initial particles | 4,070,610 | 587,351 |  | 3,701,302 | 3,376,871 |  |
| Classified Open/Closed particles | n.a./873,921 | n.a./114,159 |  | 144,067/628,561 | 803,898/227,172 |  |
| Final Closed/Locked particles | 162,468/n.a. | 84,885/20,913 |  | 395,425/n.a. | 59,857/55,580 |  |
| Symmetry imposed | C3 (Closed) | C3 (Closed) | C3 (Locked) | C3 (Closed) | C3 (Closed) | C3 (Locked) |
| Map resolution at FSC=0.143 (Å) | 3.1 | 3.5 | 3.9 | 3.1 | 3.4 | 3.2 |
| Map sharpening B-factor (Å<sup>2</sup>) | -33.1 | -24.0 | -6.7 | -48.0 | -24.4 | -22.8 |
| EMDB |  |  |  |  |  |  |
| **Refinement** |  |  |  |  |  |  |
| PDB |  |  |  |  |  |  |
| Model compositions |  |  |  |  |  |  |
| Amino acids | 3105 | 3081 | 3213 | 3105 | 3105 | 3153 |
| Glycans | 63 | 60 | 63 | 63 | 63 | 63 |
| Model *B* factors (Å<sup>2</sup>) |  |  |  |  |  |  |
| Amino acids | 125.52 | 155.40 | 154.33 | 149.64 | 139.76 | 98.39 |
| Glycans | 149.00 | 175.31 | 200.17 | 171.27 | 158.68 | 136.54 |
| R.m.s. deviations |  |  |  |  |  |  |
| Bond lengths (Å) | 0.009 | 0.007 | 0.007 | 0.007 | 0.007 | 0.004 |
| Bond angles (°) | 1.569 | 1.123 | 1.092 | 1.368 | 1.473 | 0.658 |
| Validation |  |  |  |  |  |  |
| MolProbity score | 1.53 | 1.55 | 1.85 | 1.49 | 1.48 | 1.35 |
| Clashscore | 4.50 | 4.24 | 5.22 | 3.47 | 3.45 | 2.77 |
| Rotamer outliers (%) | 0.44 | 1.12 | 1.58 | 0.33 | 1.11 | 0.66 |
| Ramachandran plot |  |  |  |  |  |  |
| Favored (%) | 95.68 | 95.62 | 93.72 | 94.83 | 95.49 | 95.89 |
| Allowed (%) | 4.32 | 3.88 | 6.19 | 4.78 | 3.93 | 4.01 |
| Disallowed (%) | 0.00 | 0.49 | 0.09 | 0.39 | 0.59 | 0.10 |

